# OvirusTdb: A Repository of Oncolytic Viruses used in Cancer Treatment

**DOI:** 10.1101/2020.01.08.896720

**Authors:** Anjali Lathwal, Rajesh Kumar, Gajendra P.S. Raghava

**Author notes:** Contributed equally to the work (Anjali Lathwal, Rajesh Kumar). <u>Mailing Address of Authors</u> Anjali Lathwal: - Rajesh Kumar: - Gajendra P.S. Raghava. Corresponding Author <u>Corresponding Author</u> Gajendra Pal Singh Raghava Head and Professor Department of Computational Biology Indraprastha Institute of Information Technology, Delhi Okhla Industrial Estate, Phase III (Near Govind Puri Metro Station) New Delhi, India - 110020 Website: http://webs.iiitd.edu.in/raghava/ Office: A-302 (R&D Block).

## Abstract

One of the emerging technologies to fight against cancer is oncolytic virus-based immunotherapy which directly lysis tumor cells. Recently, the FDA approved an oncolytic virus named T-vec for the treatment of melanoma; several hundred other viruses are in clinical trials. In order to facilitate the scientific community to fight against cancer, we build a repository of oncolytic viruses called OvirusTdb (https://webs.iiitd.edu.in/raghava/ovirustdb/). This is a manually curated repository where information is curated from research papers and patents. The current version of the repository maintains comprehensive information on therapeutically important oncolytic viruses with 5927 records where each record has 25 fields such as the virus species, cancer cell line, synergism with anti-cancer drugs, and many more. It stores information on 09 types of DNA and 15 types of RNA viruses; 300 recombinant and 09 wildtype viral strains; tested against 124 cancer types and 427 cancer cell lines. Approximately, 1047 records show improved anti-cancer response using combinatorial approach of chemotherapeutic agents with virus strains. Nearly, 3243 and 1506 records show cancer cell death via apoptosis induction and immune activation, respectively. In summary, a user-friendly web repository of oncolytic viruses for information retrieval and analysis have been developed to facilitate researchers in designing and discovering new oncolytic viruses for effective cancer treatment.

## Introduction

Cancer is the leading cause of death worldwide. According to GLOBOCAN database of world health organization (WHO) report 2018, the global cancer burden rise to 18.1 million new cases and 9.6 million deaths due to cancer [1]. This puts cancer as the second leading cause of death after cardiovascular diseases [2]. Scientists and researchers worldwide try to discover new molecules and treatment strategies to combat this deadly disease. Drugs such as paclitaxel, vincristine, vinblastine, docetaxel, NT-219 and several treatment strategies such as kinase inhibitor, mitotic disruptor, HDAC inhibitors, radiotherapy, etc. currently used to treat cancer. The current cancer treatment strategies have several limitations including increasing resistance towards chemotherapeutic agents, long anti-cancer response time, and several other cytotoxic effects [3]. An ideal anti-cancer molecule/strategy is one that selectively kills cancer cells without harming normal cells and can boosts anti-tumor immune response.

Interleukin-2 (IL) based therapy is one of the approved advance immunotherapeutic strategy for the treatment of cancer with several thousand others of its kind in clinical trials [4]. Among various immunotherapy-based strategies, oncolytic viruses (OV) are gaining much more importance as a modern age therapeutics for generating anti-cancer response [5]. Oncolytic viruses can specifically spread in tumor cells without affecting normal cells. Some oncolytic viruses such as coxsackie and measles virus have the natural tendency towards specifically targeting cancer cells while others such as adenovirus (Ad) and herpes simplex virus (HSV) are genetically modified to selectively replicate in cancer cells. Also, cancer cells have some intrinsic property which makes them more susceptible to viral infection such as resistance to apoptosis, growth suppression and defects in signalling pathways like interferon pathways (IFN). By utilizing such differences, OVs can selectively infect cancer cells and initiate lysis. Oncolysis of cancer cells further releases damage-associated molecular pattern (DAMP) and tumor-associated antigens (TAAs), which serve the basis of anti-tumor immunity generated by OV [6]. A schematic mechanism of the killing of cancer cells by the oncolytic virus is shown in Figure 1.

**Figure 1.**
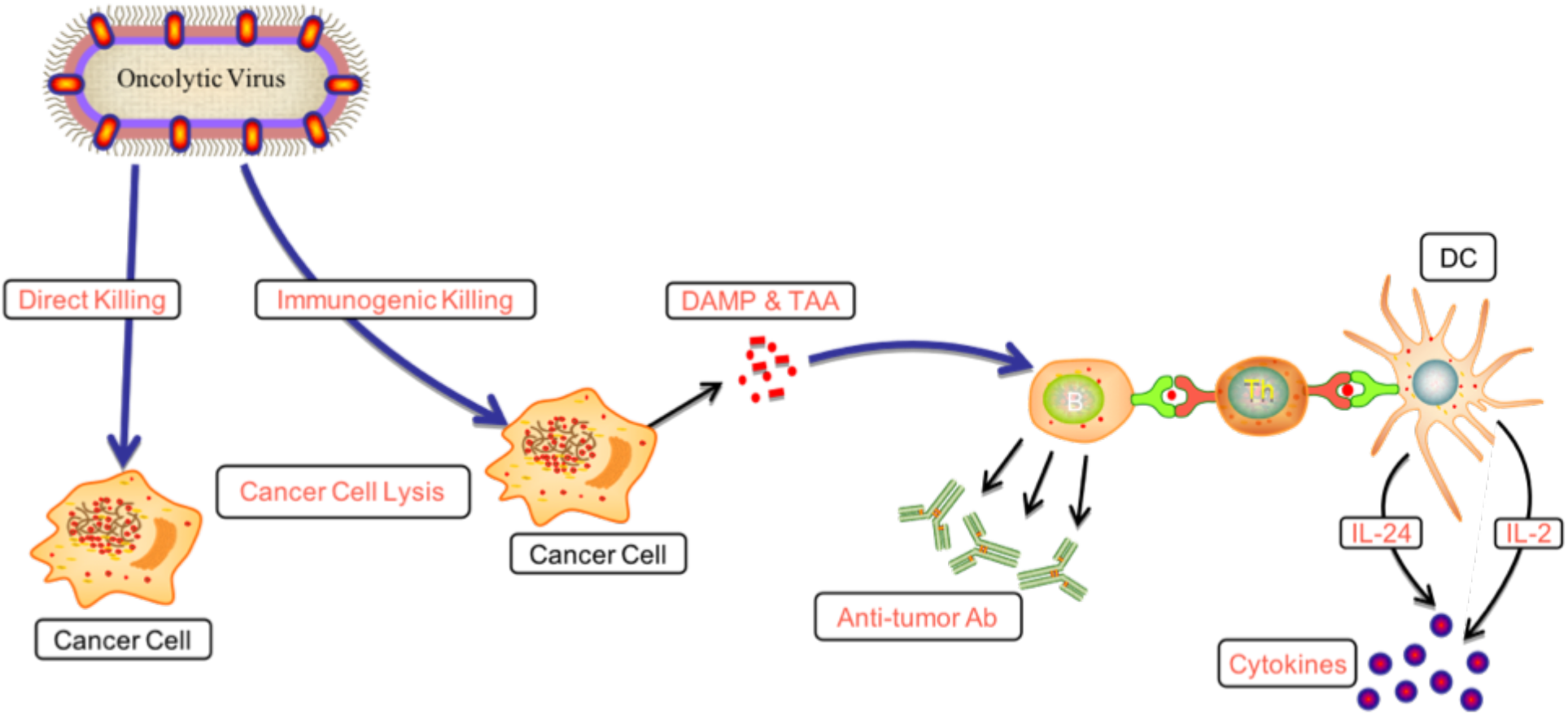
Mechanism of immunogenic cell death of cancer cells caused by the oncolytic virus. *(DAMP - Damage associated molecular pattern, TAA - Tumor-associated antigens, DC - Dendritic cells)*

Talimogene Laherparepvec (T-vec) is the only food and drug administration (FDA) approved, modified OV used for the treatment of malignant melanoma [7]. Modification in OV offers several advantages in terms of increased infectivity to a broad range of cancer cells, improved replication efficiency, and safety. For example, deletion in *ICP* and *E1B* gene of herpes simplex virus (HSV), make it to selectively replicate in cancer cells [8]. By expressing immune genes such as TNF related apoptosis-inducing ligand (TRAIL) and IL gene, one can increase the oncolytic potency of the virus in both In-vivo and In-vitro [9].

There are hundreds of OV which show promising results in various clinical and preclinical studies, but their scattered information in the literature is difficult to access. The immense therapeutic potential of OV and the gap in the literature, motivated us to make a single repository which is an homage to oncolytic viruses used to kill cancer cells. Therefore, we developed OvirusTdb (webs.iiitd.edu.in/raghava/ovirustdb), a repository that provides a manual collection of cancer-killing viruses.

## Results

### Data statistics of the OvirusTdb

OvirusTdb is a unique collection of experimentally tested oncolytic viruses collected from the literature. It contains a total of 5927 records, out of which 5456 were collected from research articles and 471 comes from patents. It provides information on 24 virus species, where the majority of viruses are modified and some are in their natural form. It also covers 124 different types of cancer and 427 different types of cancer cell lines.

One of the key advantages of using OV to treat cancer is that it also elicits an anti-tumor immune response which is also evident from the data collected as in nearly 55% of cases OV therapy induce apoptosis in the cancer cells and in 25% of cases it also triggers the immune response as seen in terms of increased interleukin secretion and T-cell activation. The complete statistics of the database are provided in Table 1. Data analysis revealed that most widely studied oncolytic virus for the treatment of cancer is the adenovirus followed by herpes simplex virus and vaccinia virus. A major portion of In-vivo studies were carried out on BALB/c mice and assay used in the majority of the studies is MTT assay as shown in Figure 2.

**Table 1.** The complete statistics of the database

| Database Statistics |  |
| --- | --- |
| Total records | 5927 (Research articles=5456 + Patents=471) |
| Total viral species | 24 (DNA genome=9 + RNA genome=15) |
| Wildtype viral strain | 15 |
| Genetically modified viral strain | 300 |
| Cancer studied | 124 |
| Cancer cell line tested | 427 |
| Assays | 30 |
| Mice/Rat species studies | 22 |
| Ex-vivo study | 1 |
| Viral strains with chemotherapeutic agents | 1047 (records) |
| Apoptosis induction by viral strains | 3243 (~55% of records) |
| Immune system activation by viral strains | 1506 (~ 25% records) |

**Figure 2.**
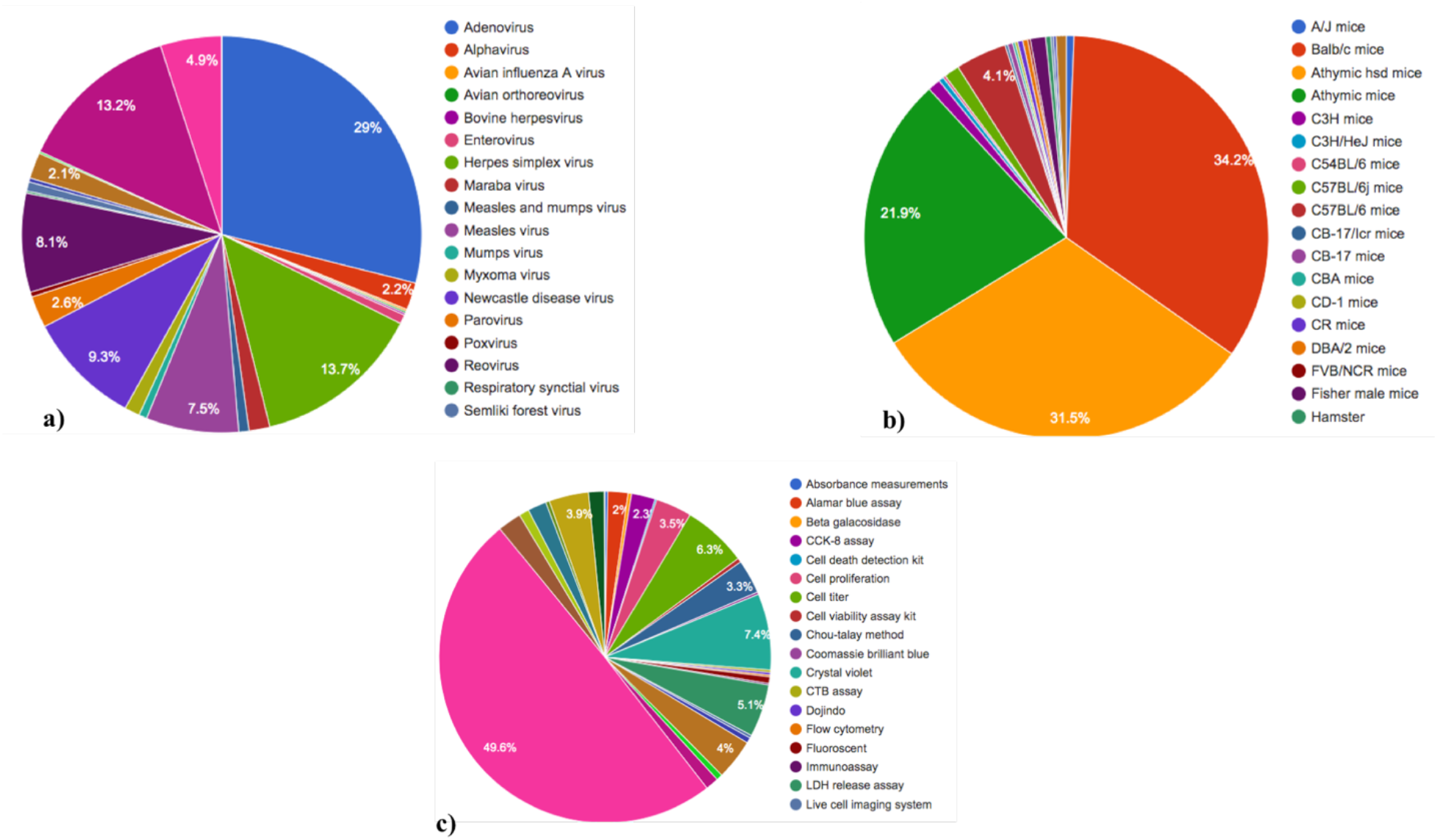
Visual representation of data statistics of: a) Oncolytic virus species used in the studies to analyse their oncolytic effects on cancer cells b) Model organism used in the studies to analyse the effect of Oncolytic viruses c) Different assays used in the studies to analyse the oncolytic effects of viruses on different cancer types.

### Utility of the database

New trends are emerging in the search for better therapy against cancer. In this regard, oncolytic viruses serve immense potential. OvirusTdb provides its users with the immense advantage of getting extensive information on oncolytic viruses at a single platform, which is otherwise difficult to access. It provides user with several facilities that includes-i) Which kind of genomic modification can improve the oncolytic potential of wild type virus, ii) What kind of combination therapy with chemotherapeutics has been used to improve the therapeutic potential of viruses, iii) What is the route of administration of virus to enhance its bioavailability, iv) Which immune and cell death pathway it activates.

### Case study: - Information retrieval and analysis on adenovirus from OvirusTdb

Here, we have shown step-by-step about what kind of information a user can get from OvirusTdb. If user is concerned about adenovirus, type adenovirus in the search box under simple search tab and click check the name box.

A number of fields are provided on the simple search page for selective retrieval of information as shown in the Figure 3. User can get all the selected available information on adenovirus stored in database by clicking on search option. For example, a total of 1716 records was found on the selected fields. Detailed information about the individual record can be obtained by clicking the respective ID. PubMed papers can be retrieved simply by clicking on the PMID which will provides additional information to the users of the database.

**Figure 3:**
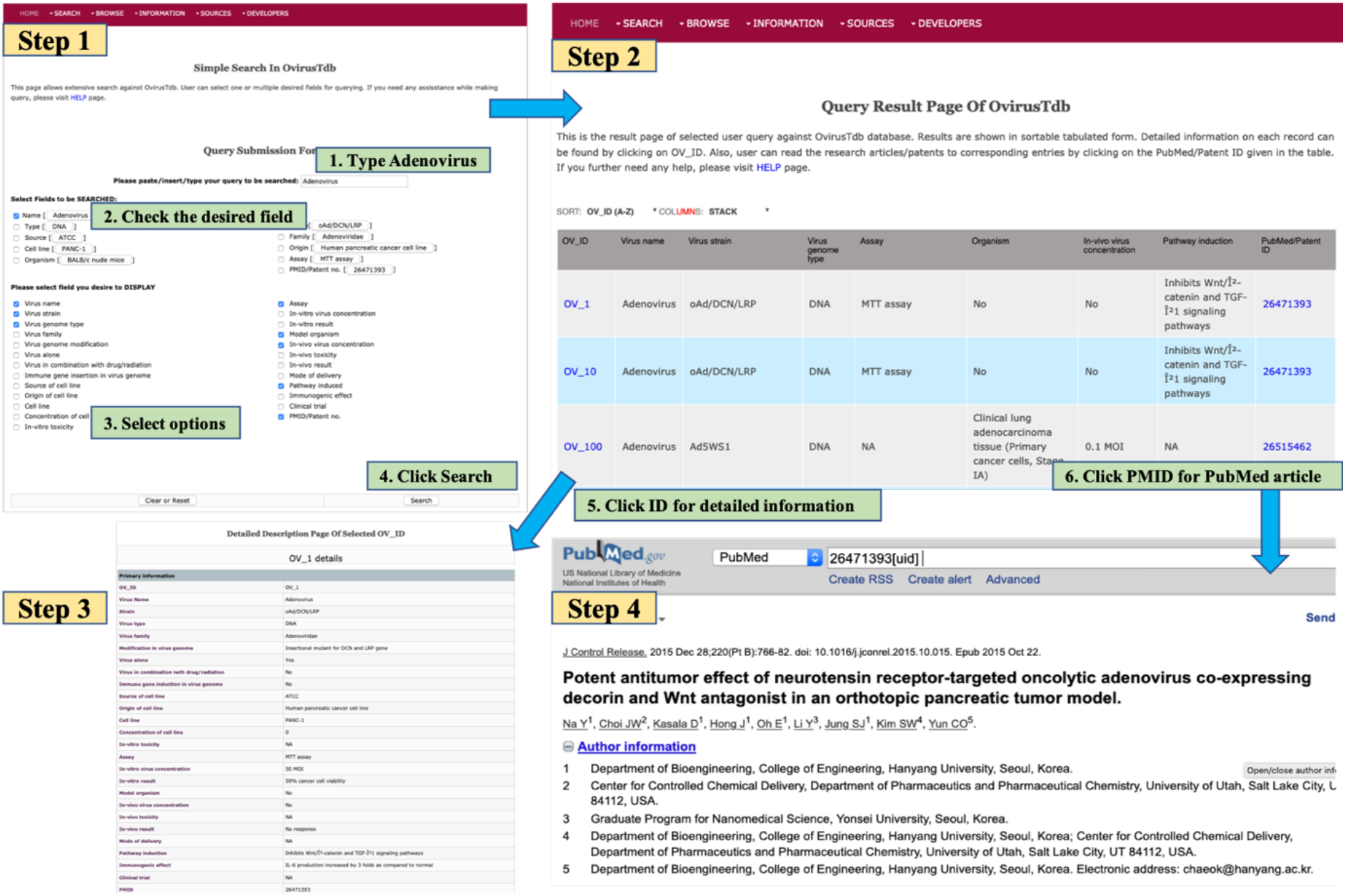
Visual representation of utility of the database

### Data submission, limitation and update of OvirusTdb

We aim to make OvirusTdb - a single platform of the available information related to the oncolytic viruses used in cancer therapeutics. The online data submission platform under sources drop down menu enables the user of the database to submit information in OvirusTdb. Nonetheless, curators of the database will ensure the legitimacy of the new entry before including it in the OvirusTdb. The data stored in OvirusTdb was manually curated and thoroughly checked to mitigate the mistake, but it would be unfair to claim absolute consistency due to human mistakes that may have arisen. In order to maintain the consistency, we will update the OvirusTdb, preferably after every three years.

## Discussion

Viruses that have the ability to infect and kill cancer cells are termed as “Oncolytic viruses”. Some viruses have the natural ability to infect cancer cells while some others are genetically modified to show their desired oncolytic effect. The importance of using OV to treat cancer is seen in terms of that not only does it kills cancer cells but also generate an anti-tumor immune response, which inhibits the remission after withdrawal of therapy.

The therapeutic potential of oncolytic viruses has been experimentally tested in various animal and tumor models including dozens of OV in clinical trials to treat human malignancies [10]. There are several challenges associated with OV therapy where one such challenge is the replication of virus and its clearance from the system as host immune cells are competent enough to control virus replication [4]. Another major factor that needs to be considered for successful OV therapy is the induction of the right immune response. Several studies are trying to elucidate the activation of right DAMP which can induce the right cytokine to induce immunogenic cell death (ICD) [11,12] of cancer.

Despite such limitation, T-vec is the only OV, approved by the FDA for the treatment of skin melanoma. T-vec is a genetically modified type −1 HSV currently being evaluated in several countries for combination therapy with chemo and radiotherapy agents [13].

Despite such a huge therapeutic application, there is no such database that collectively provides information on all the experimentally tested oncolytic viruses present in literature to treat cancer. Therefore, it is necessary to catalogue all the information available in the literature regarding OV. Thus, we have developed a database “OvirusTdb” which provides information on OV with total 5924 records. To the best of author’s knowledge, there is no such database present in literature which covers almost all the aspects of virus-based cancer therapy. We believe that this database is useful for virologists, immunologists, oncologists and biotechnologists who wish to design and use oncolytic viruses with/without combination approach for the treatment of cancer.

Although the strategy of using OV to treat cancer seems to be fascinating, still there is much more to be explored in this area [14]. The approval of T-vec by the FDA further promoted and strengthened the research in oncolytic virus based therapy (OVT). There is need for critical monitoring of patients treated with OV for the presence of immune biomarkers, type of cancer cell death induced by DAMP to understand how OVT works in population.

### Comparison with other available resources on viruses

There are many resources available on the viruses such as VIPERdb – maintain information on 816 viral capsid structures, viruSITE – provides all genomic information on viruses and viroid published in NCBI, ViFi - a tool for detecting viral integration and fusion mRNA sequences from Next Generation Sequencing data, and ViPR – catalogue information on immune epitopes, host factor and antiviral drugs for 19 virus families. None of the above mentioned available resources holds experimentally validated and detailed information on the oncolytic potential of viruses for cancer treatment. OvirusTdb unlike other resources provides user with comprehensive information on 24 viral species and their therapeutic efficacy on 427 cancer cell lines, including the information on any immune and apoptotic pathways it may activates.

We believe that if information from basic and clinical research is implemented in a proper way, the oncolytic virus based therapeutics may serve as a powerful approach in the battle against cancer.

## Materials and Methods

### Data acquisition

All the data on oncolytic viruses were collected from the research articles and patents. The relevant articles were collected from PubMed using a combination of the keywords “Oncolytic virus” and “oncolytic virotherapy” up to May 2019. A total of 4514 articles were found in PubMed using the above-mentioned search terms. All the articles were manually screened for relevant experimental details and 1604 articles were filtered. Patents were searched from United States Patents and Trademark Office (USPTO) site using the above-mentioned keywords, results in 644 patents. All the articles and patents manually screened for relevant information and final data comes from 166 research articles and 27 patents.

### Database architecture and web interface

All the information available in the literature related to OV was stored in the SQL table and presented in the form of a user-friendly and interactive web interface named OvirusTdb. OvirusTdb is built on Linux based Apache server (LAMP). The responsive front-end web interface of OvirusTdb was developed using bootstrap, a popular responsive development framework including HTML, CSS and JavaScript and MySQL queries were used to manage the back-end data. The complete architecture of the OvirusTdb is presented in Figure 4.

**Figure 4.**
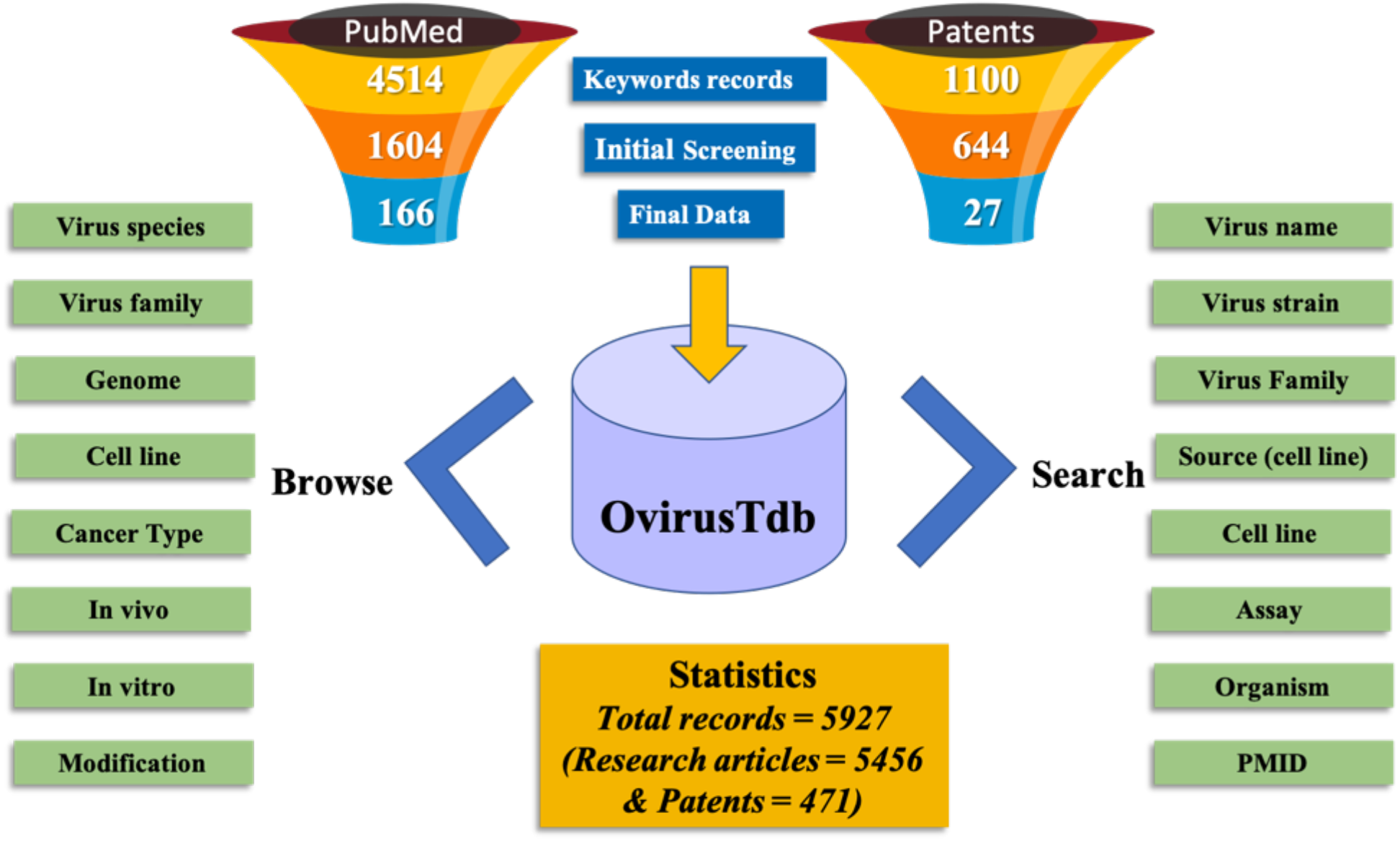
Database organization and architecture of OvirusTdb

### Primary information

The manually collected information from research articles and patents is provided in tabular form under different fields in the database. This primary information can be broadly categorized into 1) Virus information-represents name, strain, genome, family and any modification of virus, 2) In-vitro information-represents cell line, cell line source, assay used to measure virus killing efficiency, 3) In-vivo information - represents animal model used and killing efficiency of virus, 4) Immune booster information - includes after killing which pathway is activated namely caspases and apoptosis, and generation of secondary immune response in terms of interleukins and T-cells.

### Searching

This facility allows the users to search within the database in a time-efficient manner using a query against any field of databases such as name, strain, type, cytotoxicity, cell line, PMID, and the patent number. This simple search tool allows users to customize their search criteria by selecting the desired fields. Advanced search criteria provide users the additional facility of multiple querying using numerous fields available for selection.

### Browsing

A user-friendly browsing facility is provided on the website to retrieve information from the database in an effortless manner. Users can browse on major fields which include-1) Name of virus species 2) cancer type 3) Cell line source 4) Model organism 5) Assay information.

## Authors’ contributions

G.P.S.R. made substantial contributions to the conception and design of the work, data analysis, supervision, funding acquisition, project administration and drafting of the manuscript. A.L. and R.K. made substantial contributions to data collection, manual curation, web interface development, data analysis, and manuscript preparation.

## Competing interest

The authors declare no conflict of interest.

## Acknowledgments

Authors are thankful to funding agencies University Grant Commission (UGC) and Council of Scientific and Industrial Research (CSIR), Indraprastha Institute of Information Technology, New Delhi (IIIT-D), and Govt. of India for financial support and fellowships.

## Funding

This work was supported by J. C. Bose Fellowship (Grant number SRP076), Department of Science and Technology, India.

## Abbreviations

WHO: World Health Organization
IL: Interleukin
OV: Oncolytic Virus
Ad: Adenovirus
HSV: Herpes Simplex Virus
IFN: interferon
DAMP: Damage Associated Molecular Patterns
TAA: Tumor-Associated Antigens
T-VEC: Talimogene Laherparepvec
TRAIL: TNF related apoptosis-inducing ligand
USPTO: United States Patents and Trademark Office
LAMP: Linux based Apache server
OVT: Oncolytic Virus Therapy
ICD: Immunogenic Cell Death

## References

[1] R.L. Siegel, K.D. Miller, A. Jemal, Cancer statistics, 2019, CA. Cancer J. Clin. 69 (2019) 7–34. https://doi.org/10.3322/caac.21551.

[2] H. Nagai, Y.H. Kim, Cancer prevention from the perspective of global cancer burden patterns., J. Thorac. Dis. 9 (2017) 448–451. https://doi.org/10.21037/jtd.2017.02.75.

[3] V. Hanusova, L. Skalova, V. Kralova, P. Matouskova, Potential anti-cancer drugs commonly used for other indications., Curr. Cancer Drug Targets. 15 (2015) 35–52. https://doi.org/10.2174/1568009615666141229152812.

[4] M.E. Davola, K.L. Mossman, Oncolytic viruses: how “lytic” must they be for therapeutic efficacy?, Oncoimmunology. 8 (2019) e1581528. https://doi.org/10.1080/2162402X.2019.1596006.

[5] H.L. Kaufman, F.J. Kohlhapp, A. Zloza, Oncolytic viruses: a new class of immunotherapy drugs., Nat. Rev. Drug Discov. 14 (2015) 642–62. https://doi.org/10.1038/nrd4663.

[6] K.A. Schalper, H.S. Kim, J. Raja, S.N. Gettinger, J.M. Ludwig, Oncolytic virus immunotherapy: future prospects for oncology, J. Immunother. Cancer. 6 (2018) 140. https://doi.org/10.1186/s40425-018-0458-z.

[7] P.K. Bommareddy, A. Patel, S. Hossain, H.L. Kaufman, Talimogene Laherparepvec (T-VEC) and Other Oncolytic Viruses for the Treatment of Melanoma., Am. J. Clin. Dermatol. 18 (2017) 1–15. https://doi.org/10.1007/s40257-016-0238-9.

[8] J.L. Hummel, E. Safroneeva, K.L. Mossman, The role of ICP0-Null HSV-1 and interferon signaling defects in the effective treatment of breast adenocarcinoma., Mol. Ther. 12 (2005) 1101–10. https://doi.org/10.1016/j.ymthe.2005.07.533.

[9] P. Sova, X.W. Ren, S. Ni, K.M. Bernt, J. Mi, N. Kiviat, A. Lieber, A tumor-targeted and conditionally replicating oncolytic adenovirus vector expressing TRAIL for treatment of liver metastases, Mol. Ther. 9 (2004) 496–509. https://doi.org/10.1016/j.ymthe.2003.12.008.

[10] V. Cervera-Carrascon, R. Havunen, A. Hemminki, Oncolytic adenoviruses: a game changer approach in the battle between cancer and the immune system., Expert Opin. Biol. Ther. 19 (2019) 443–455. https://doi.org/10.1080/14712598.2019.1595582.

[11] A.D. Garg, S. More, N. Rufo, O. Mece, M.L. Sassano, P. Agostinis, L. Zitvogel, G. Kroemer, L. Galluzzi, Trial watch: Immunogenic cell death induction by anticancer chemotherapeutics., Oncoimmunology. 6 (2017) e1386829. https://doi.org/10.1080/2162402X.2017.1386829.

[12] A. Showalter, A. Limaye, J.L. Oyer, R. Igarashi, C. Kittipatarin, A.J. Copik, A.R. Khaled, Cytokines in immunogenic cell death: Applications for cancer immunotherapy., Cytokine. 97 (2017) 123–132. https://doi.org/10.1016/j.cyto.2017.05.024.

[13] R. Kanai, H. Wakimoto, T. Cheema, S.D. Rabkin, Oncolytic herpes simplex virus vectors and chemotherapy: are combinatorial strategies more effective for cancer?, Future Oncol. 6 (2010) 619–34. https://doi.org/10.2217/fon.10.18.

[14] S. Farkona, E.P. Diamandis, I.M. Blasutig, Cancer immunotherapy: The beginning of the end of cancer?, BMC Med. 14 (2016). https://doi.org/10.1186/s12916-016-0623-5.

